# MZF1-mediated GAPDH overexpression drives glycolytic reprogramming and neuroendocrine progression in advanced prostate cancer

**DOI:** 10.64898/2026.06.03.729867

**Authors:** Wang Liu, Lily He, Cuncong Zhong, Yuzhuo Wang, Da Zhang, Moben Mirza, Benyi Li

**Author notes:** **Corresponding author**: Benyi Li, MD/PhD, KUMC Urology.

## Abstract

Drug resistance to the androgen receptor (AR) antagonist is a critical obstacle in the clinic for advanced prostate cancers. Especially, AR antagonist treatment-induced neuroendocrine progression represents a lethal and therapy-resistant subtype. Although transcriptional and epigenetic lineage plasticity have been extensively implicated in treatment-induced neuroendocrine progression, the contribution of metabolic adaptation remains incompletely understood. Here, we identified a previously unrecognized metabolic reprogramming mechanism induced by AR antagonists in castration-resistant prostate cancer (CRPC) models. AR antagonist treatment markedly enhanced glycolytic activity and induced glyceraldehyde-3-phosphate dehydrogenase (GAPDH) expression. Genetic depletion of GAPDH suppressed AR antagonist-induced glycolytic activation, altered transcriptomic and metabolic programs, reduced neuroendocrine-associated marker expression, and inhibited xenograft tumor growth. Mechanistically, GAPDH promoter pulldown coupled with mass spectrometry, siRNA screening, and chromatin immunoprecipitation assays identified myeloid zinc finger-1 (MZF1) as a key transcription factor for Enzalutamide-induced GAPDH gene expression. Pharmacological inhibition of GAPDH using koningic acid (KA) or penta-O-galloyl-β-D-glucopyranose (PGG) significantly suppressed tumor growth and attenuated neuroendocrine-associated molecular programs in CRPC cell-derived xenograft and patient-derived t-NEPC xenograft models. Collectively, our findings identify an AR antagonist-induced MZF1-GAPDH signaling axis that promotes glycolytic activation and neuroendocrine-associated metabolic adaptation during treatment resistance. These results support targeting GAPDH-dependent metabolic reprogramming as a potential therapeutic strategy for treatment-resistant prostate cancer.

**Graphic abstract:** 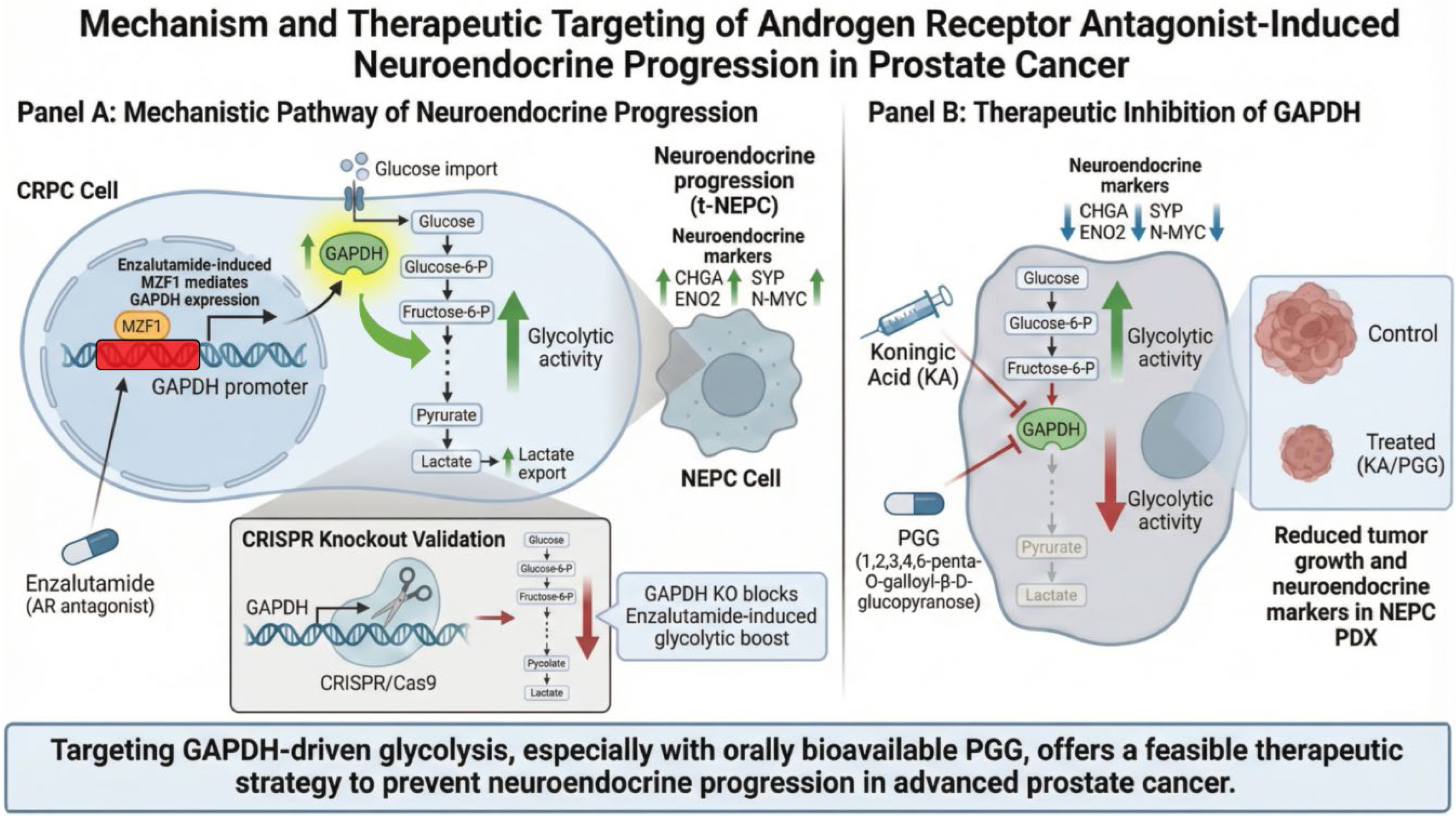

## Introduction

Metastatic prostate cancers were primarily treated with the androgen deprivation therapy (ADT); however, patients inevitably develop castration resistance, emerging as castration resistant prostate cancer (CRPC) (1). Since the androgen receptor (AR) signaling pathway is active in the CRPC cells, AR antagonists are widely used as the first-line treatment in CRPC patients (2). With the widespread use of potent AR antagonist drugs, including Enzalutamide and Abiraterone, about 20-25% of CRPC patients develop neuroendocrine trans-differentiation (so-called treatment-induced NEPC, or t-NEPC) (3), accounting for more than 25-30% mortality without means to cure in the clinic (4, 5). In the last two decades, a large body of studies showed multiple oncogenes, epigenetic modulators, transcription factors, RNA splicing factors, and even long non-coding RNAs that were involved in NEPC progression (reviewed in refs. (3–10)), reflecting the mechanistic complexity.

Metabolic reprogramming in malignant cells is characterized by the deregulation of metabolic pathways for the needs of rapid tumor growth and cancer cell proliferation (11). These altered metabolic pathways include increased uptake of glucose and diverging metabolic intermediates from glycolysis/Krebs cycle to biosynthesis (11). Among these metabolic reprogrammings, increased aerobic glycolysis is one of the main features in human malignancies (12), including t-NEPC (13, 14). Our recent studies with patient-derived xenografts (PDX) revealed that the t-NEPC model LTL-331R exerted a highly upregulated glycolytic activity (15), one of the malignant hallmarks featured in metabolic reprogramming (11). In fact, targeting the altered glycolysis pathway in cancer cells has emerged as a potent cancer therapy (15, 16). Especially, inhibition of glyceraldehyde-3-phosphate dehydrogenase (GAPDH), a critical glycolytic enzyme, achieved a profound anti-cancer outcome specifically in highly glycolytic cancers (17–19). This evidence indicates that the overactivated GAPDH-dependent glycolytic activity in the highly glycolytic tumors, such as the t-NEPC, might provide a feasibility for therapeutic intervention (17).

GAPDH is a homo-tetramer that mainly resides in the cytoplasmic compartment and acts as the essential component of cellular glycolytic machinery (12). In addition, GAPDH also acts as a quintessential moonlight protein with different activities in multiple cellular compartments (20). In the 6th step of glycolytic flux, GAPDH utilizes inorganic phosphate to produce ATP-generating organic intermediates, a critical step for cellular energy production (21). Recent studies suggest that GAPDH may function as an important flux-controlling enzyme during aerobic glycolysis, particularly in highly glycolytic cancer cells (22).

Although the GAPDH gene is considered a housekeeping gene and its expression level at the mRNA or protein levels was widely used as an internal control for gene expression studies, its aberrant expression was reported in multiple cancers associated with poor prognosis, including prostate cancers (23–25). Early studies showed that GAPDH expression was significantly increased in Dunning R-3327 rat prostate cancer cells compared to normal prostate tissue and that increased GAPDH expression was closely associated with cancer cell motility (26). In normal human prostate tissues, GAPDH protein was abundantly detectable in the nuclei of basal and stromal cells but not in secretory epithelial cells (27). Conversely, prostate cancer tissues from late-stage patients exerted a strong cytoplasmic staining of GAPDH protein and increased GAPDH expression, which correlated with tumor grade and metastasis (23). All these data provide a strong rationale for GAPDH as a drug target of anti-glycolysis therapy (25).

In an effort to illustrate the mechanistic insight for overactivated glycolytic activities during NEPC progression after AR antagonist treatment (15), and also to develop a novel treatment for t-NEPC patients, we investigated the glycolytic pathway in CRPC and NEPC tissues. Our investigation was focused on the GAPDH gene expression and functional alterations during t-NEPC progression. Our results showed that AR antagonist treatment largely increased GAPDH expression *via* the transcription factor myeloid zinc finger-1 (MZF1)-dependent mechanism and strongly enhanced glycolytic activities. Suppressing GAPDH activity with an orally available natural compound, 1,2,3,4,6-penta-O-galloyl-β-D-glucopyranose (PGG), and a well-known compound, koningic acid (KA), completely blocked neuroendocrine xenograft tumor growth and diminished neuroendocrine features in NEPC PDX models. Therefore, targeting the GAPDH-dependent glycolytic pathway represents a feasible therapeutic intervention for NEPC patients.

## Materials and methods

### Cell culture, special chemical reagents, plasmid constructs, and antibodies

Prostate cancer NCI-H660 (ATCC CRL-5813; RRID: CVCL_1576), PC-3 (ATCC CRL-1435; RRID: CVCL_0035), 22Rv1 (ATCC CRL-2505; RRID: CVCL_1045), C4-2B (ATCC CRL-3315/3314; RRID: CVCL_4784), LNCaP (ATCC CRL-1740; RRID: CVCL_0395), and benign prostate BPH1 (RRID: CVCL_1091) cell lines were obtained from ATCC (Manassas, VA) as described (28). Cell lines were authenticated by Short Tandem Repeat (STR) profiling within the last 6 months. PC-3, 22Rv1, C4-2B, and BPH1 cells were cultured in RPMI 1640 media containing 10% fetal bovine serum (FBS) and 1% penicillin/ streptomycin at 37°C in a humidified atmosphere containing 5% CO_2_. The NCI-H660 cell line was cultured in RPMI 1640 medium containing 5% FBS, insulin/transferrin/selenium (1:100), hydrocortisone (10 nM), beta-estradiol (10 nM), and L-glutamine (2 mM). Cell lines were routinely tested for mycoplasma contamination and confirmed to be negative. LTL352 and LTL331R NEPC PDX models were obtained from the Living Tumor Lab to generate subcutaneous transplant xenografts in NOD-SCID-Gamma2 (NSG) mice (Jackson Labs). The reagents and compounds for this study are listed in Supplemental Table S6. The antibodies for this study were listed in Supplemental Table S7.

### Glycolytic activity assays

Cells were treated under the indicated conditions and cultured in RPMI-1640 medium supplemented with 2% charcoal-filtered fetal bovine serum (cFBS) for hormone-deprivation experiments. Glucose uptake was measured using the Glucose Uptake-Glo™ Assay (Promega, J1341) or the Glucose Uptake Cell-Based Assay Kit (Cayman Chemical, No. 600470). Glucose consumption was determined using the Glucose (GO) Assay Kit (Sigma-Aldrich, GAGO20-1KT) and calculated as [(initial glucose − residual glucose)/initial glucose] × 100%. L-lactate production was quantified using the L-Lactate Assay Kit (Cayman Chemical, No. 700510). GAPDH enzymatic activity was measured using the GAPDH Activity Colorimetric Assay Kit (Abcam, ab204732). All assays were performed according to the manufacturers’ instructions, and values were normalized to total cell number or protein content as indicated.

### Western blot, Cellular thermal shift assay (CETSA), peptide probe labeling, and fluorescent gel analysis

Cells were washed twice and harvested with cold PBS solution, then lysed with Radioimmunoprecipitation assay (RIPA) lysis buffer with protease/phosphatase inhibitor cocktail (Cell Signaling Technologies, Danvers, MA, USA). Protein concentrations were determined with the Standard BCA Protein Assay Kit (Thermo Fisher, Waltham, MA, USA). Equal amounts of protein were resolved on 6-15% SDS-PAGE gels and transferred onto PVDF membranes. Proteins were transferred to the PVDF membrane. The membranes were blocked for 60 min with 5% no-fat milk solutions prepared in Tris Buffered Saline (TBS), incubated overnight at 4°C with the indicated dilution of the primary antibodies. Appropriate peroxidase-conjugated secondary antibody (1:5000 dilution) was used. Protein bands were visualized using an ECL solution from Santa Cruz Biotech (Santa Cruz, CA, USA).

Enzalutamide binding with the MZF1 proteins was examined using the CETSA assay, as described previously (29). Briefly, C4-2B cells were treated with Enzalutamide (10 μM) for 2 h. Cell pellets were washed with PBS, followed by two repeated freeze-thaw cycles with liquid nitrogen. The lysates were then aliquoted and heated at increasing temperatures (37–69°C) for 3 min. and then cooled down on ice for 2 min. The cell lysates were briefly vortexed and then centrifuged at 18,000 g for 20 min at 4°C. The supernatant was loaded onto an SDS-PAGE gel for western blot analysis.

To determine GAPDH activity with the SEC-6 fluorescent probe (30), C4-2B cells were treated with Enzalutamide (5 μM) for the indicated durations. Cell lysates (50 μl, 3 mg/mL) in PBS were incubated with the SEC6 probe (100 μM) at RT for 1 h. In-gel fluorescent signals were visualized on a Bio-Rad ChemiDoc MP imaging system. After fluorescent visualization, gels underwent a typical procedure for Coomassie staining and destaining. Stained gels were visualized on a Bio-Rad ChemiDoc MP imaging system.

### Quantitative real-time PCR (qRT-PCR)

Total RNA samples were isolated using TRIzol RNA isolation reagents. The RNA samples were treated with DNase to remove genomic DNA contamination. Reverse transcription and real-time PCR reactions were conducted according to the manufacturer’s instructions. Primer sequences are listed in Supplementary Table S8. Quantitative PCR data were analyzed using the 2^-ΔΔCt^ method. The relative mRNA expression levels of target genes (CHGA, SYP, MYCN, ENO2, and GAPDH) were normalized to the endogenous control 18S rRNA.

### GAPDH promoter pulldown assay and MZF1 identification

A DNA-protein pulldown assay was established to identify transcription factors responsible for AR antagonist-induced GAPDH upregulation. The 0.5KB (-488/+21 upstream of the transcription start site) GAPDH promoter fragment was excised by XhoI/EcoRI double digestion from the hGAPDH-rLUC construct (Addgene #82479). The promoter fragment was gel-purified and labeled with biotin using a DNA 3’-End labeling kit (ThermoFisher Scientific, catalog #89818) (Fig S2A). C4-2B cells grown in 15 cm culture dishes were switched to medium containing 2% charcoal-filtered FBS (cFBS) overnight and treated with Enzalutamide (10 µM) or DMSO for 1 h. Cell lysates were prepared for the DNA-binding assay. The gel-purified biotin-tagged fragments were then mixed with C4-2B cell lysate at 4 °C for 4 hours, in the presence of poly (dI-dC) to minimize non-specific interactions. DNA-bound proteins were pulled down using the Streptavidin beads. After high-stringency washing, the bound protein complexes were eluted and subjected to in-gel tryptic digestion followed by Liquid Chromatography-Tandem Mass Spectrometry (LC-MS/MS) for comprehensive proteomic profiling. Data were searched against the UniProt human database using MaxQuant software (version 2.0.3, RRID: SCR_014485).

### RNA-seq assay and data processing

Raw RNA-seq samples were preprocessed with Trimmomatic v0.38 (31), and then aligned to the human genome (32), indexed with NCBI RefSeq (33), and all annotations with STAR (version 2.7.0d, RRID: SCR_004463) (34). For downstream analysis, raw read quantities were fed to DESeq2 (35) to compute the differential expression level between any two groups, and the RPKM-normalized quantities were transformed to z-score and visualized using a heatmap, which was hierarchically clustered by Pearson’s correlation. Differentially expressed genes were defined using an adjusted p-value <0.05 and absolute log2 fold change >1. The genes were clustered and associated with metabolic pathways using the comprehensive DAVID Knowledgebase (36). For upregulated genes (log_2_FC > 2), analysis included Gene Ontology (GO) enrichment, pathway network connectivity, and hierarchical pathway clustering. Gene set enrichment analysis (GSEA) was further conducted against the MSigDB hallmark gene sets to identify coordinated transcriptional programs. For downregulated genes (log_2_FC < -2), GO enrichment and pathway network connectivity were analyzed to identify suppressed biological modules. Enrichment significance was determined using a modified Fisher’s exact test; all p-values were adjusted for multiple testing using the Benjamini-Hochberg procedure to maintain a False Discovery Rate (FDR) < 0.05.

### Gas Chromatography-Mass Spectrometry (GC-MS) Analysis

Cellular or animal plasma metabolites were extracted in cold methanol, dried under nitrogen, and derivatized with methoxyamine hydrochloride in pyridine, followed by N-methyl-N-(trimethylsilyl) trifluoro-acetamide (BSTFA). Metabolomic profiling was performed by gas chromatography-mass spectrometry (GC-MS) at the University of Utah Metabolomics Core Facility. Metabolites were separated on a 30 m × 0.25 mm × 0.25 μm DB-5ms capillary column with helium carrier gas at 1.0 mL/min. Samples (1.0 μL) were injected in splitless mode at 250°C. The GC oven was held at 70°C for 1 min, ramped to 300°C at 10°C/min, and held for 5 min. The mass spectrometer was operated in electron-impact ionization mode (70 eV) with a scan range of m/z 50-600. Metabolites were identified by comparison to reference spectra, and peak areas were normalized to total protein content measured before extraction.

Metabolomic data were acquired and processed using Agilent MassHunter software. Metabolite identification was performed by matching retention times and mass fragmentation patterns against a combination of an in-house library (validated with pure standards), the NIST library, and the Fiehn library. Peak areas were integrated using MassHunter Quant and normalized for comparative analysis. Statistical analyses of metabolomic data were performed using the web-based Metaboanalyst 5.0, as described in our previous publication (30).

### Chromatin immunoprecipitation (ChIP)

C4-2B cells grown in 15 cm culture dishes were switched to medium containing 2% charcoal-filtered FBS (cFBS) overnight and treated with Enzalutamide (10 µM) or DMSO for 1 h. Chromatin immunoprecipitation (ChIP) was performed using the SimpleChIP® Plus Sonication Chromatin IP Kit (#56383, Cell Signaling Technology) according to the manufacturer’s instructions. Briefly, C4-2B cells were cross-linked with 1% formaldehyde and quenched with glycine. Chromatin was fragmented to 150-900 bp by sonication. Equal amounts of chromatin were incubated with anti-MZF1 antibody (Santa Cruz Biotechnology, sc-293218;1:50) or Normal Rabbit IgG (CST#2729) overnight at 4°C, followed by capture with ChIP-Grade Protein G Magnetic Beads. After proteinase K digestion and DNA elution using spin columns, the enrichment of MZF1 at the GAPDH promoter was quantified by qPCR and normalized to 2% input DNA. The Human RPL30 gene (CST#7015) was used as an internal control for normalization.

### CRISPR/CAS9 knockout (KO) system stable cell line

GAPDH knockout (KO) stable cell line was established in C4-2B cells using the CRISPR plasmid system (sc-420485) obtained from Santa Cruz (Santa Cruz, CA, USA). The protocol provided by Santa Cruz Biotech was followed to generate the GAPDH/KO cell lines. Stable subline cells with GAPDH reinstallation were established with the wild-type or mutant GAPDH plasmids in C4-2B cells.

### Xenograft and PDX animal experiments

Prostate cancer cell lines (C4-2B, 22Rv1, and NCI-H660) were cultured in RPMI-1640 medium containing 10% FBS and grown overnight. The cells were harvested during the logarithmic growth phase, washed twice with sterile PBS, and then resuspended at a concentration of 5×10^7^ cells/ml in sterile PBS. The cell suspensions were subcutaneously injected into 6-week-old nu/nu nude mice. Male mice (6 weeks old; n=8/group) were randomly assigned to treatment groups using a random number generator. All animal procedures were performed in accordance with institutional guidelines and approved by the Institutional Animal Care and Use Committee (IACUC). Animal welfare was monitored daily.

LTL352 and LTL331R were established as previously reported (37). After a quick thaw in a 37°C water bath, the tissue pieces were washed once with sterile Hank’s Balanced Salt Solution (HBSS) and then kept on ice. Implantation in NOD-SCID-Gamma2 (NSG) mice (Jackson Labs) was conducted immediately. Once the xenograft tumors were palpable, animals were castrated, followed by different treatments for 3 weeks.

### Immunohistochemistry (IHC)

Formalin-fixed paraffin-embedded (FFPE) sections (4 μm) from CRPC and NEPC xenograft tissues were deparaffinized in xylene and rehydrated through graded ethanol solutions to distilled water. Antigen retrieval was performed in citrate buffer (10 mM sodium citrate, pH 6.0) by heat-induced epitope retrieval for 15–20 min, followed by cooling to room temperature. Endogenous peroxidase activity was quenched using 3% hydrogen peroxide for 10 min, and nonspecific binding was blocked using 5% bovine serum albumin (BSA) for 30 min at room temperature.

Sections were incubated overnight at 4°C with primary antibodies against GAPDH, ENO2, SYP, and Ki67 at the indicated dilutions. Following PBS washing, sections were incubated with secondary antibodies using the VECTASTAIN Elite ABC Universal PLUS Peroxidase Kit (Vector Laboratories, PK-8200, Newark, CA, USA) according to the manufacturer’s instructions. Immunoreactivity was visualized using 3,3′-diaminobenzidine (DAB) chromogen, followed by hematoxylin counterstaining. Sections were subsequently dehydrated through graded ethanol solutions, cleared in xylene, mounted, and imaged by light microscopy.

Staining intensity and the percentage of positively stained cells were assessed independently by investigators blinded to sample identity. Quantification was performed using an H-score or by calculating the percentage of positive cells, where indicated.

### Statistical analysis

Data are presented as mean ± SEM from at least three independent experiments unless otherwise stated. Comparisons between two groups were performed using two-tailed Student’s t-tests, whereas comparisons among multiple groups were analyzed using one-way ANOVA followed by post hoc testing. Statistical analyses were performed using GraphPad Prism v9.0 (RRID: SCR_002798), and p < 0.05 was considered statistically significant.

## Results

### Enzalutamide treatment enhanced glycolytic activities in CRPC cells

To determine whether androgen receptor (AR) antagonists alter cellular metabolism, we first examined glycolytic changes induced by Enzalutamide under three culture conditions: serum-supplemented medium (10% FBS), serum-free medium, and hormone-deprived medium containing 2% charcoal-stripped FBS (cFBS), the latter serving as an *in vitro* model of androgen deprivation therapy (ADT). As expected, glucose consumption in C4-2B cells was markedly reduced under serum-free or ADT conditions compared with serum-supplemented conditions (Fig 1A). Interestingly, Enzalutamide significantly increased glucose consumption only under hormone-deprived conditions, whereas no significant effects were observed in serum-supplemented or serum-free conditions (Fig 1A).

**Fig 1.**
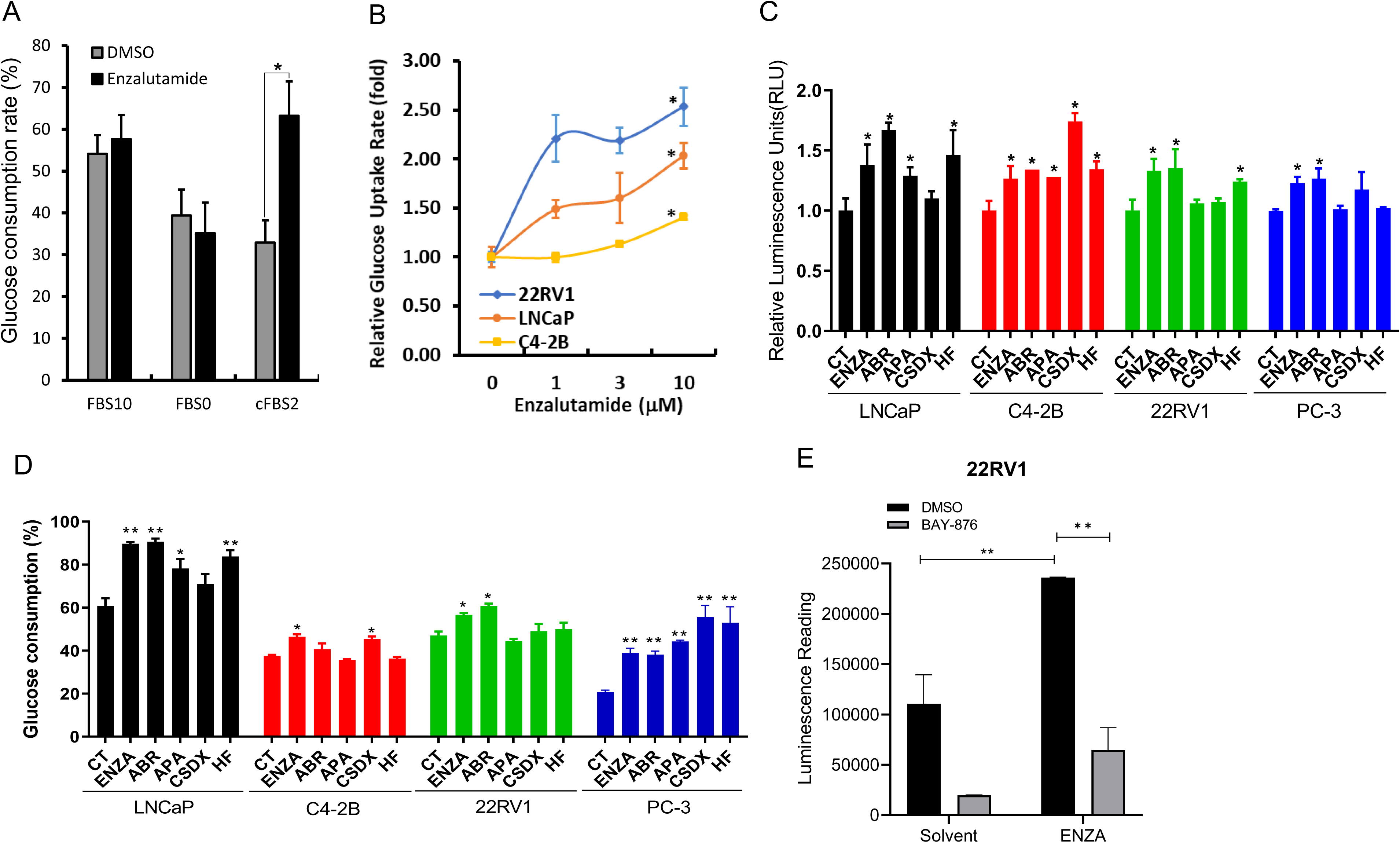
AR antagonist enhances glycolytic activity in CRPC cells. **A** C4-2B cells were treated with DMSO or Enzalutamide (10 µM) for 24 h under 10% FBS (FBS10), serum-free (FBS0), or 2% cFBS conditions. Glucose levels in cell media were measured after treatment. Consumption rate was calculated as: [(before-after)/before x 100%]. **B** LNCaP, C4-2B and 22Rv1 cells were treated with DMSO or Enzalutamide at indicated concentrations for 4 h in 2% cFBS conditions, followed by glucose uptake analysis. **C** Cells were treated with DMSO or the AR antagonists as indicated for 24 h under 2% cFBS conditions. Glucose uptake was assessed at the treatment using the bioluminescent Glucose Uptake-Glo™ Assay kit (Promega). **D** Cells were treated with DMSO or the AR antagonists as indicated for 24 h in 2% cFBS condition. Glucose consumption was measured using a glucose oxidase-based assay. **E** 22Rv1 cells were pre-treated with glucose transporter inhibitor BAY-876 (1 µM), followed by the solvent or Enzalutamide (ENZA, 10 µM) for 24 h under 2% cFBS conditions. Glucose uptake was then measured. Quantitative data were presented as mean ± SEM. The asterisks indicate statistical significance compared to the solvent or DMSO control (ANOVA analysis; * p < 0.05, ** p < 0.01).

We then measured glucose uptake activity. Under ADT conditions, Enzalutamide induced a concentration-dependent increase of glucose uptake in AR-positive LNCaP, C4-2B, and 22Rv1 cells (Fig 1B). Interestingly, Enzalutamide also increased glucose uptake in AR-negative PC-3 cells (Fig 1C), whereas no significant effect was observed in benign prostate epithelial BPH1 cells (Fig S1A). Consistent with these findings, additional AR antagonists, including Abiraterone (ABI), Apalutamide (APA), Bicalutamide (Casodex, CSDX), and Hydroxy-Flutamide (HF), also increased glucose uptake to varying degrees in LNCaP, C4-2B, 22Rv1, and PC-3 cells (Fig 1B).

Consistent with the glucose uptake results, AR antagonists also increased glucose consumption to various extents in LNCaP, C4-2B, 22Rv1, and PC-3 cells (Fig 1D). Pretreatment with the glucose transporter inhibitor BAY-876 abolished Enzalutamide-induced glucose uptake (Fig 1E), indicating a requirement for glucose transporter activity in this response. Collectively, these findings suggest that AR antagonists trigger adaptive glycolytic reprogramming under ADT conditions through non-canonical stress-associated signaling mechanisms.

### AR antagonist treatment increases GAPDH expression in CRPC cells

Given the central role of GAPDH in glycolytic metabolism and previous findings implicating GAPDH in cancer metabolism (17), we next investigated whether Enzalutamide regulates GAPDH expression. Enzalutamide treatment in C4-2B, 22Rv1, and PC-3 cells largely increased GAPDH protein levels, whereas no significant changes were observed in BPH1 cells (Fig S1B). The induction of GAPDH protein expression occurred in both a dose-and time-dependent manner (Fig 2A-2B). Consistent with these observations at the protein level, Enzalutamide treatment also significantly increased GAPDH mRNA levels under ADT conditions (Fig 2C), which was in a dose-dependent manner (Fig 2D).

**Fig 2.**
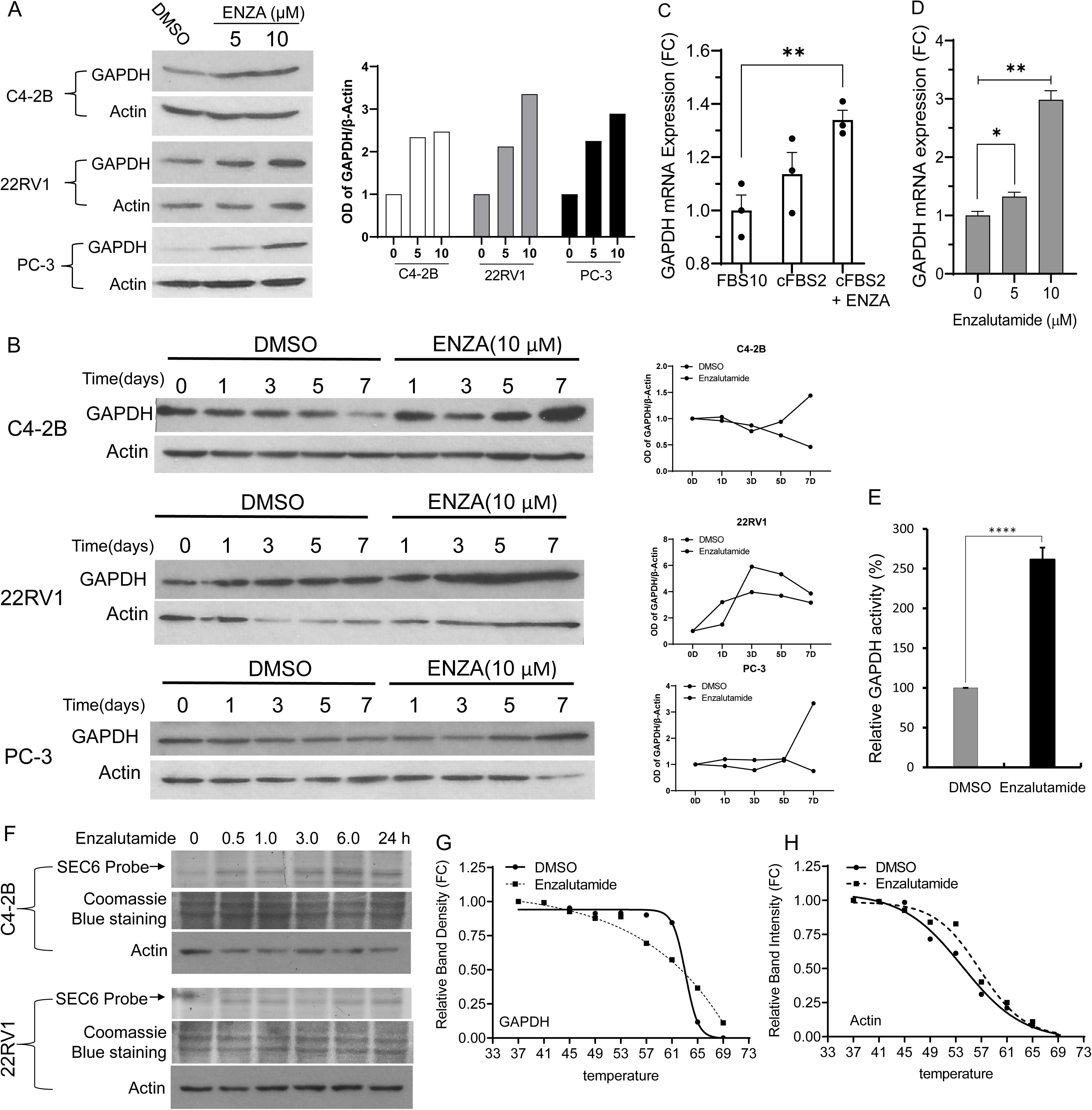
Enzalutamide induces GAPDH expression and enzymatic activity in CRPC cells. **A** Cells were treated with DMSO or Enzalutamide at the indicated concentrations for 5 days under 2% cFBS conditions, followed by immunoblot analysis. **B** Cells were treated with DMSO or Enzalutamide (10 µM) under 2% cFBS conditions for the indicated time, followed by GAPDH protein analysis. **C, D** Cells were treated with DMSO or Enzalutamide (10 μM) under the indicated culture conditions for 24 h. Total RNA was extracted, and GAPDH mRNA expression was measured by qRT-PCR. **E** C4-2B Cells were treated with or without Enzalutamide in 2% cFBS conditions for 24 h. GAPDH enzymatic activity was quantified in cell lysates using a colorimetric assay kit (Abcam). **F** Cells were treated with Enzalutamide (10 μM) for the indicated times. Cell lysates were incubated with the SEC6 probe (100 μM) for 1 h at room temperature, followed by SDS-PAGE and fluorescence imaging. Coomassie blue staining and Actin immunoblotting served as loading controls. **G, H** C4-2B cells were treated with DMSO or Enzalutamide (10 µM) for 1 h. Cells were harvested for the CETSA assay as described (72). Data are presented as mean ± SEM. The asterisks indicate statistical significance compared with the solvent or parental control (ANOVA analysis; ** *p* < 0.01, **** *p* < 0.0001).

To determine whether increased GAPDH expression translated into functional changes, GAPDH enzymatic activity was assessed using both a cell-based GAPDH activity assay and a fluorescent probe-based assay. Enzalutamide treatment significantly increased GAPDH activity in both assays (Fig 2E-2F) (38). We then examined if Enzalutamide interacted with the GAPDH protein in a CETSA assay. As shown in Fig 2G, Enzalutamide treatment did not dramatically cause GAPDH melting curve shift, similar to the internal control protein Actin (Fig 2H). Collectively, these data demonstrated that Enzalutamide treatment enhanced GAPDH gene expression at the mRNA level and subsequently protein accumulation, leading to glycolytic overactivation.

### GAPDH activity is required for Enzalutamide-induced glycolytic overactivation

To verify the GAPDH contribution to Enzalutamide-induced glycolytic overactivation, GAPDH gene knockout C4-2B subline cells were generated using the CRISPR/Cas9 strategy. Another C4-2B subline with a stable re-installation of wild-type GAPDH gene was also established (Fig 3A). GAPDH gene knockout markedly reduced basal glucose uptake (Fig 3B) and glucose consumption (Fig 3C), accompanied by a decrease in glycolytic activity as assessed by L-lactate production (Fig 3D). To further determine the role of GAPDH in AR antagonist-induced metabolic changes, GAPDH-deficient cells were treated with Enzalutamide or Abiraterone. GAPDH depletion substantially attenuated AR antagonist-induced glucose consumption (Fig 3E) and L-lactic acid production (Fig 3F). These data demonstrated the involvement of GAPDH activity in AR antagonist-induced glycolysis.

**Fig 3.**
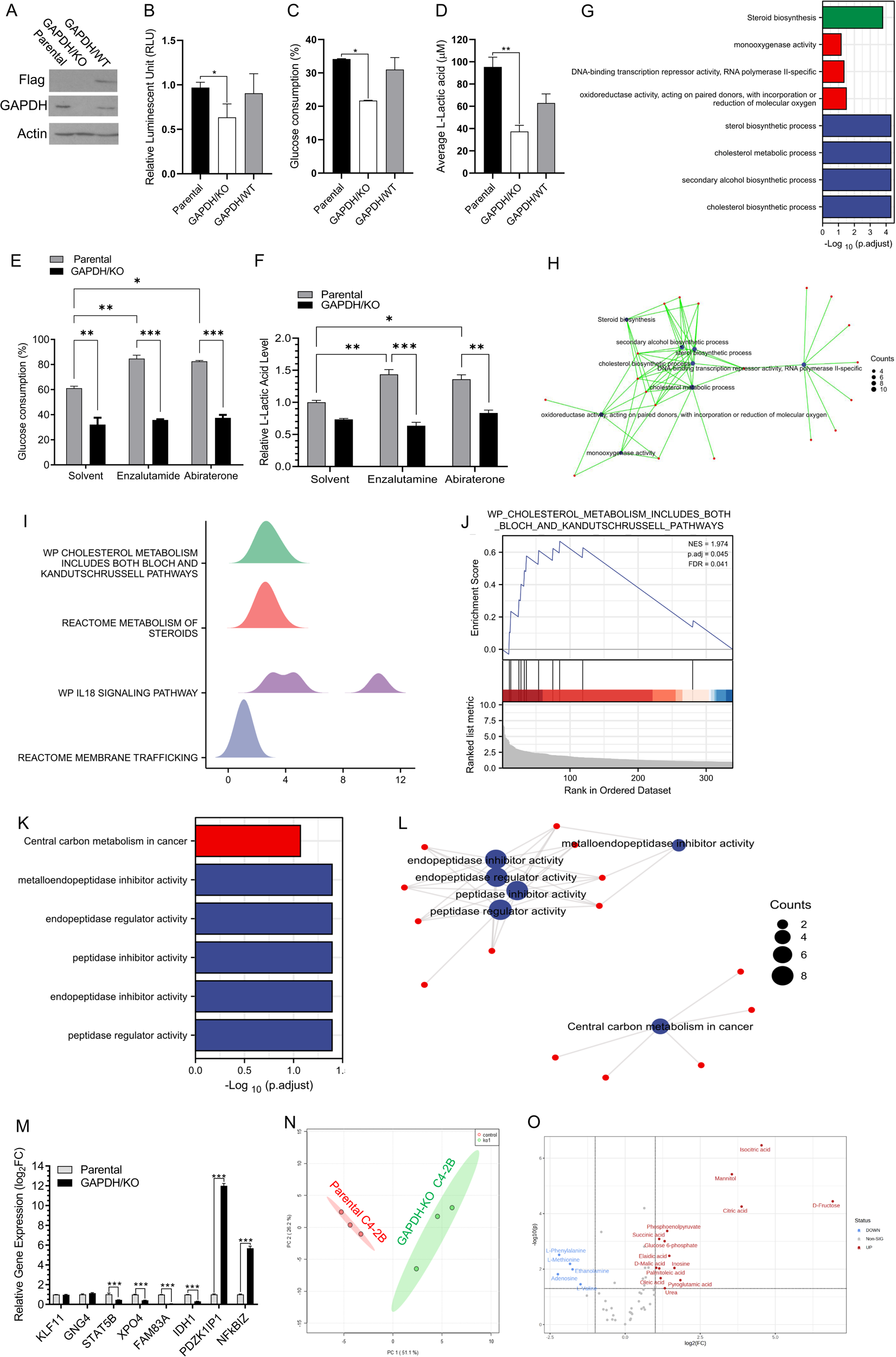
GAPDH depletion suppresses glycolytic activity and induces transcriptomic and metabolic remodeling in CRPC cells. **A** GAPDH was depleted in C4-2B cells using a CRISPR/Cas9-mediated HDR strategy. GAPDH-deficient sublines were further rescued by stable expression of Flag-tagged wild-type GAPDH. Actin served as a loading control. **B** Glucose uptake was measured in parental, GAPDH-knockout, and GAPDH-rescued C4-2B cells. **C, D** Cells were cultured overnight, and fresh medium was added for 24 h. Glucose consumption and L-lactate production were measured and normalized to total cell number. **E, F** Parental and GAPDH-knockout C4-2B cells were treated with vehicle, Enzalutamide (10 μM), or Abiraterone (10 μM) for 24 h. Glucose consumption and L-lactate production were measured. **G-L** Whole-transcriptome RNA-seq analysis was conducted using parental and GAPDH/KO subline C4-2B cells. Gene Ontology (G), pathway network connection (H), pathway cluster (I), and Gene Set Enrichment Analysis (J) were performed for the upregulated genes (log2FC > 2). In addition, gene ontology (K) and pathway network connection (L) were performed for downregulated genes (log_2_FC < -2). **M** Selected differentially expressed genes were validated by qRT-PCR in parental and GAPDH-knockout C4-2B cells. **N, O** Metabolites were extracted from exponentially growing parental and GAPDH-knockout C4-2B cells and analyzed by GC-MS. Principal component analysis was used to compare global metabolic profiles (N), and volcano plot analysis was used to visualize differential metabolite changes (O). Data are presented as mean ± SEM. Asterisks indicate statistical significance compared with the solvent control (ANOVA analysis; * *p* < 0.05, ** *p* < 0.01, *** *p* < 0.001).

### GAPDH depletion induces transcriptomic and metabolic reprogramming in CRPC cells

To investigate the broader consequences of GAPDH depletion, transcriptomic profiling was performed using the RNA sequencing (RNA-seq) approach. GAPDH knockout resulted in significant transcriptional alterations, with 276 genes upregulated by more than two-fold compared with parental cells (Supplemental Table S1). Gene Ontology analysis revealed that most of these increased genes are related to cholesterol biosynthesis and transcriptional repressors (Fig 3G-3H). Consistent with these findings, GSEA analysis revealed four major pathways, including the cholesterol metabolism genes (Fig 3I-3J, Supplemental Table S2). In parallel, GAPDH knockout downregulated 389 genes by more than two-fold (Supplemental Table S1). Most of these downregulated genes were related to central carbon metabolism and peptidase inhibitor activity (Fig 3K-3L). GSEA further identified six major pathways related to peptidase activity (Table 1).

**Table 1.** Pathway enrichment for GAPDH-KO downregulated genes.

To validate these transcriptomic findings, eight genes with drastic alterations after GAPDH knockout were verified in a real-time qPCR assay. Consistent with RNA-seq results, the upregulation of *PDZK1IP*1 and *NFKBIZ*, as well as the downregulation of *STAT5B, XPO4, FAM83A,* and *IDH1,* was confirmed exponentially (Fig 3M). Unexpectedly, *KLF11* and *GNG4* expression were not validated.

The metabolic effect of GAPDH knockout was assessed with a metabolomic GC-MS approach (Supplemental Table S3). Principal component analysis (PCA) revealed a drastic difference between the C4-2B parental and GAPDH-KO cells (Fig 3N). Glycolytic metabolite D-fructose and Tricarboxylic acid (TCA) cycle metabolite citric acid were significantly accumulated in GAPDH-KO cells, and five amino acids were reduced following GAPDH depletion (Fig 3O, Table 2). These data suggest that GAPDH knockout caused substantial changes in cellular energy production, including glycolytic and TCA cycle, and amino acid metabolism.

**Table 2.** Major alterations of GC-MS metabolite analysis.

### GAPDH depletion suppresses CRPC xenograft growth

We next evaluated the impact of GAPDH depletion on tumor growth *in vivo*. Parental C4-2B cells, GAPDH-knockout C4-2B cells, and GAPDH-reinstated C4-2B cells with wild-type GAPDH were implanted subcutaneously into castrated nude mice to establish xenograft tumors. As shown in Fig. 4A, tumors derived from GAPDH-knockout C4-2B cells exhibited a markedly reduced growth compared with tumors generated from parental cells or GAPDH-rescued cells. Consistently, both tumor weight and final tumor burden were significantly reduced in the GAPDH-knockout group relative to the parental and GAPDH-rescued groups (Fig 4B, C). No significant changes in body weight were observed during treatment (Fig 4D). These data indicate that GAPDH activity is essential for CRPC xenograft growth *in vivo*.

**Fig 4.**
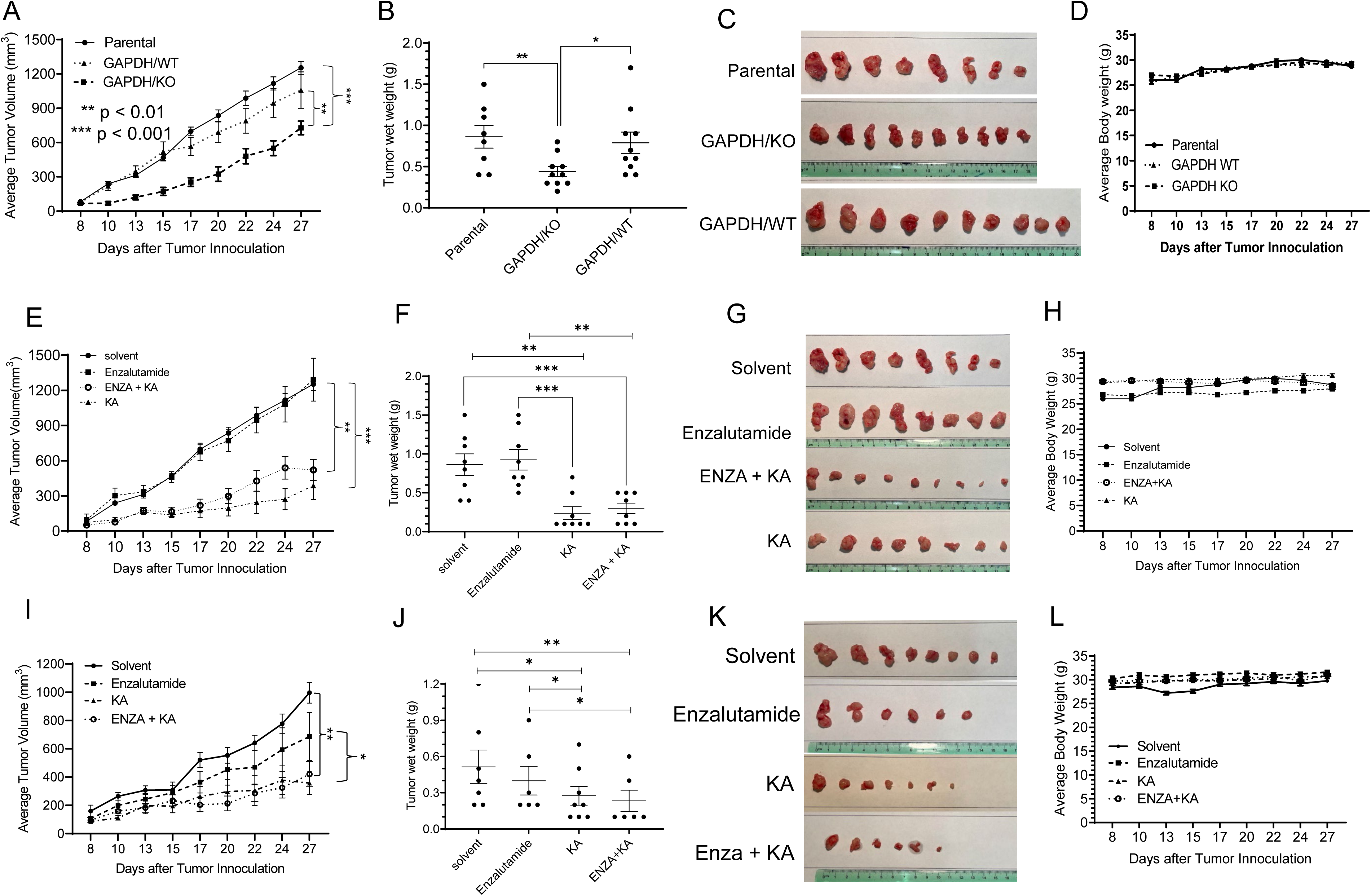
GAPDH depletion or inhibition suppresses CRPC xenograft growth. **A-D** Subcutaneous xenografts were established in castrated male nude mice using parental C4-2B cells, GAPDH-knockout cells, or GAPDH-rescued cells expressing wild-type GAPDH. Tumor volume was measured three times per week (A). Tumor weights and representative tumor images were recorded at endpoint (B, C), Body weight was monitored during treatment (D). **E-H** C4-2B cell-derived xenografts were treated with vehicle, Enzalutamide (30 mg/kg), KA (1.0 mg/kg), or Enzalutamide plus KA in castrated male nude mice. Tumor volume was measured three times per week (E). Tumor weights and representative tumor images were recorded at endpoint (F, G). Body weight was monitored during treatment (H). **I-L** 22Rv1 cell-derived xenografts were treated using the same regimen as described for C4-2B xenografts. Tumor volume (I), endpoint tumor weight (J), representative tumor images (K), and body weight were analyzed (L). Data are presented as mean ± SEM. Asterisks indicate statistical significance compared with the control group by two-way ANOVA; *p < 0.05, **p < 0.01, ***p < 0.001.

### GAPDH inhibition overcomes anti-AR resistance in CRPC xenograft models

We next evaluated the therapeutic potential of targeting GAPDH in CRPC xenograft models exhibiting resistance to AR antagonists. A previously reported GAPDH inhibitor, KA (19), was administered intraperitoneally at 1.0 mg/kg based on prior studies (39). C4-2B and 22Rv1 cell-derived xenograft models were treated with vehicle, Enzalutamide, KA, or the combination regimen. As expected, Enzalutamide treatment alone failed to suppress tumor growth in C4-2B xenografts (Fig 4E – 4G). In parallel, GAPDH expression was increased following Enzalutamide treatment, consistent with the observations in cultured cells (Fig S1C, D). In contrast, treatment with KA alone or in combination with Enzalutamide significantly inhibited tumor growth (Fig 4E – 4G). No significant changes in body weight were observed during treatment (Fig 4H), suggesting acceptable tolerability.

To further validate these findings, the same treatment regimen was evaluated in 22Rv1 cells-derived xenografts. Although Enzalutamide alone produced a modest growth inhibition, treatment with KA alone or in combination with Enzalutamide significantly reduced tumor growth (Fig 4I-4K). Similarly, no significant body weight changes were observed during treatment (Fig 4L). Collectively, these findings suggest that pharmacological inhibition of GAPDH activity suppresses CRPC tumor growth and overcomes resistance to AR-targeted therapy.

### GAPDH inhibition suppresses NEPC xenograft tumor growth

We next evaluated the antitumor effect of the GAPDH inhibition in NEPC models. NCI-H660 cells were used to establish subcutaneous NEPC xenograft tumors in nude mice. Animals were randomly assigned to receive vehicle or KA. As shown in Fig 5A-5C, KA treatment significantly suppressed tumor growth at 1.0 mg/kg without any obvious weight loss (Fig 5D).

**Fig 5.**
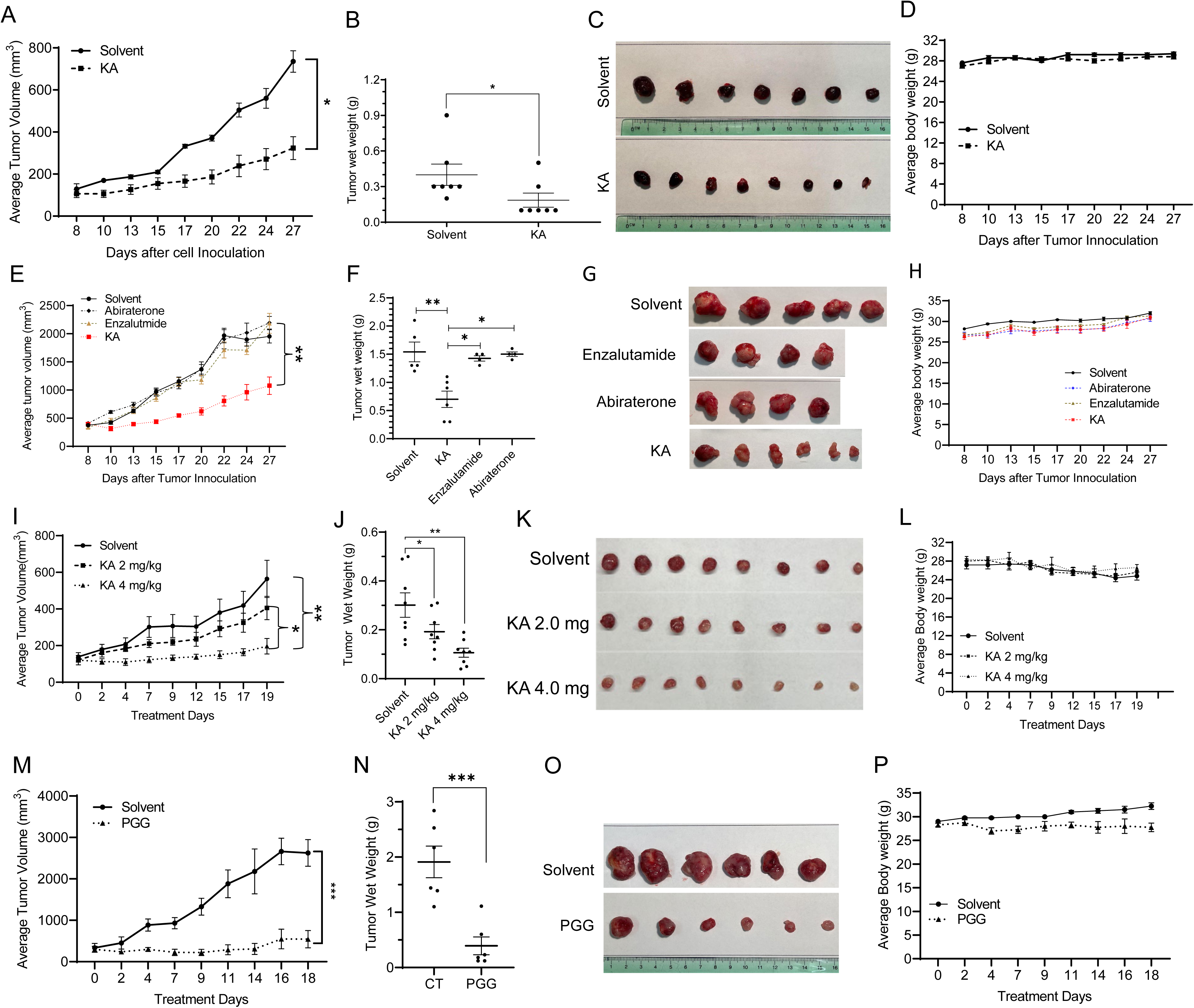
GAPDH inhibition suppresses NEPC xenograft and PDX tumor growth. **A-D** NCI-H660 cells were used to establish subcutaneous NEPC xenografts in castrated male nude mice. Animals were randomly assigned to receive vehicle or KA (1.0 mg/kg). Tumor volume was measured three times per week (A). Tumor weights and representative tumor images were recorded at endpoint (B, C). Body weight was monitored during treatment (D). **E-H** LTL331R PDX models were subcutaneously established in castrated male NSG mice and treated with Enzalutamide (30 mg/kg) or KA ( 1.0 mg/kg). Tumor growth rate was monitored using the caliper measurement three days a week (E). Tumor wet weights and mass images were recorded at the end of treatment after harvest (F, G). Animal body weights were recorded at each treatment (H). **I-L** LTL352 PDX models were subcutaneously established in castrated male NSG mice. Animals were randomly assigned to three groups to receive KA (2.0 or 4.0 mg/kg) for three weeks. Tumor growth rate was monitored using the caliper measurement three days a week (I). Tumor wet weights and mass images were recorded at the end of treatment after harvest (J, K). Animal body weights were recorded at each treatment (L). **M-P** LTL331R PDX models were subcutaneously established in castrated male NSG mice and treated with PGG (60 mg/kg) for three weeks. Tumor growth rate was monitored using the caliper measurement three times a week (M). Tumor wet weights and mass images were recorded at the end of treatment after harvest (N, O). Animal body weights were recorded at each treatment (P). Data are presented as mean ± SEM. Asterisks indicate statistical significance compared with the vehicle group by two-way ANOVA; * *p* < 0.05, ** *p* < 0.01.

To assess the therapeutic efficacy of GAPDH inhibition in clinically relevant models, two treatment-emergent NEPC (t-NEPC) patient-derived xenograft (PDX) models, LTL331R and LTL352, were evaluated in NSG mice. Consistent with resistance to AR-targeted therapy, as shown in Fig 5E-5G), neither Enzalutamide (30 mg/kg) nor Abiraterone (100 mg/kg) affected tumor growth in the LTL331R model. In contrast, KA treatment at 1.0 mg/kg significantly inhibited tumor growth without affecting body weight (Fig 5H). Interestingly, in the LTL352 PDX model, significant tumor growth inhibition was observed only at a higher KA dose (4.0 mg/kg) (Fig 5I-5L), suggesting a moderate sensitivity to KA in this model.

Lastly, we tested a natural compound, PGG, which was recently shown as a reversible GAPDH inhibitor (40). Oral administration of PGG at 60 mg/kg markedly suppressed LTL331R tumor growth in NSG mice (Fig 5M-5O), although a slight reduction (without statistical significance) in body weight was observed during treatment (Fig 5P).

### GAPDH inhibition suppresses neuroendocrine marker expression

We next investigated whether GAPDH inhibition affects the expression of neuroendocrine-associated markers in CRPC and NEPC models. Previous studies reported that hormone deprivation combined with Enzalutamide treatment markedly increased the expression of multiple neuroendocrine markers, including CHGA, SYP, ENO2, and N-MYC in C4-2B cells (3). In our study, GAPDH knockout substantially reduced the expression of these proteins, although SYP exhibited a comparatively less reduction (Fig 6A). Consistent with these findings, Enzalutamide-induced expression of neuroendocrine markers was markedly reduced following KA treatment in LTL331R-derived PDX tissues (Fig 6B).

**Fig 6.**
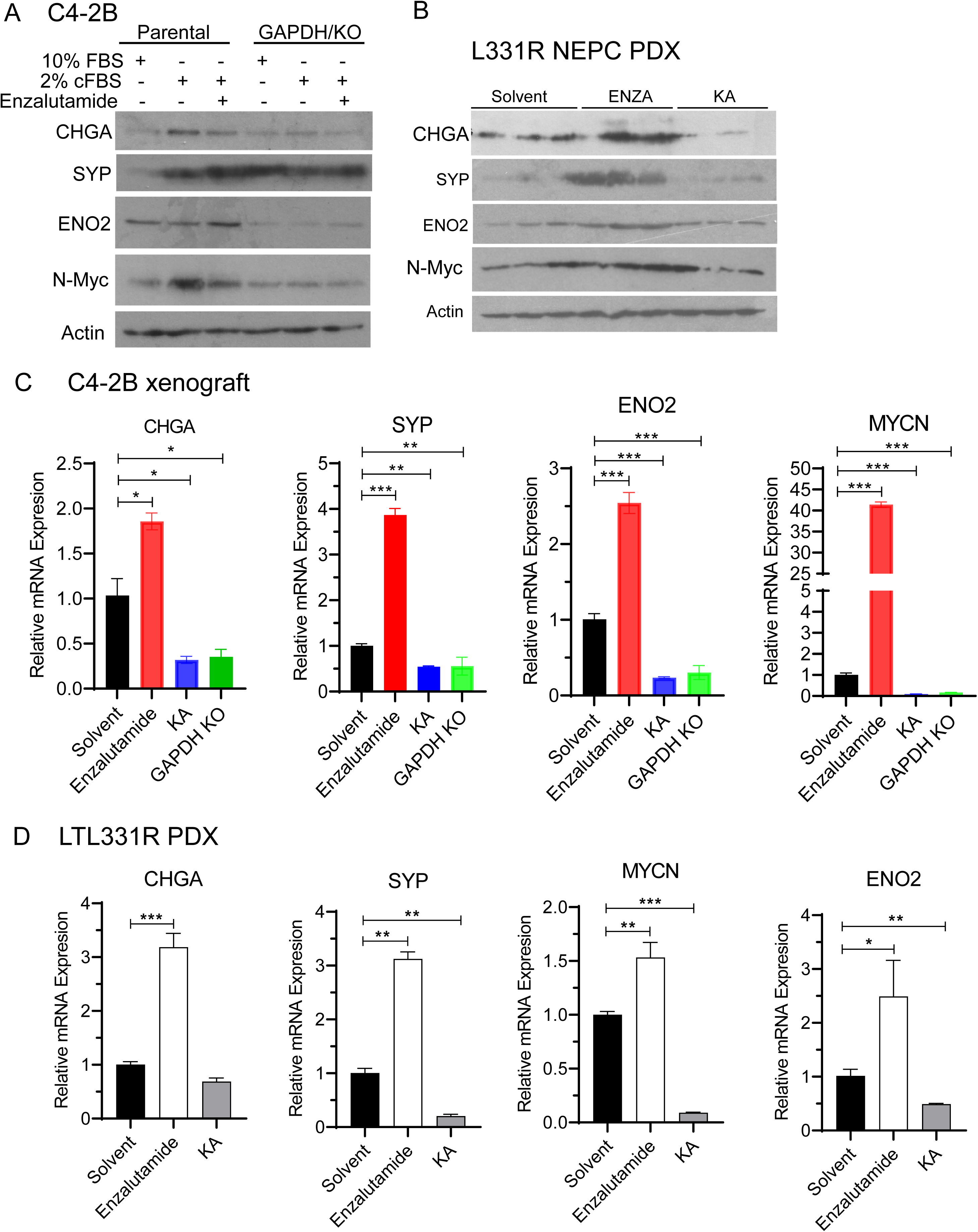
GAPDH inhibition suppresses neuroendocrine-associated molecular programs. **A** Parental and GAPDH/KO C4-2B cells were treated with or without Enzalutamide (10 μM) for 24 h under the indicated serum conditions. NEPC biomarker proteins were analyzed by immunoblotting. **B** Protein lysates from LTL331R PDX tissues were analyzed by immunoblotting using the indicated antibodies. Actin served as a loading control. **C, D** Total RNA was extracted from C4-2B cell-derived xenografts treated with Enzalutamide, KA, or GAPDH knockout, as well as from treated NEPC PDX tumors. Neuroendocrine-associated gene expression was analyzed by qRT-PCR.

At the mRNA level, the expression of neuroendocrine-associated genes was significantly elevated in Enzalutamide-treated C4-2B xenografts and LTL331R PDX tissues (Fig 6C, D). GAPDH inhibition with KA or GAPDH gene knockout eliminated their expression (Fig 6C, D). These data confirmed that GAPDH inhibition by KA and genetic depletion significantly reduced the expression of these neuroendocrine-associated genes that were strongly induced following Enzalutamide treatment.

At the protein level, we conducted an immunohistochemical analysis to further demonstrate the alterations induced by GAPDH inhibition. Our results showed that ENO2 and SYP expression were significantly reduced following GAPDH inhibition or GAPDH depletion compared with control tumors in all xenograft and PDX models (Fig 7), consistent with immunoblot and qRT-PCR results. In addition, KA treatment significantly reduced Ki67-positive cells in xenograft tumors (Fig 8). These results demonstrated that GAPDH inhibition suppressed NEPC-related gene expression at the mRNA and protein levels, leading to reduced cell proliferation.

**Fig 7.**
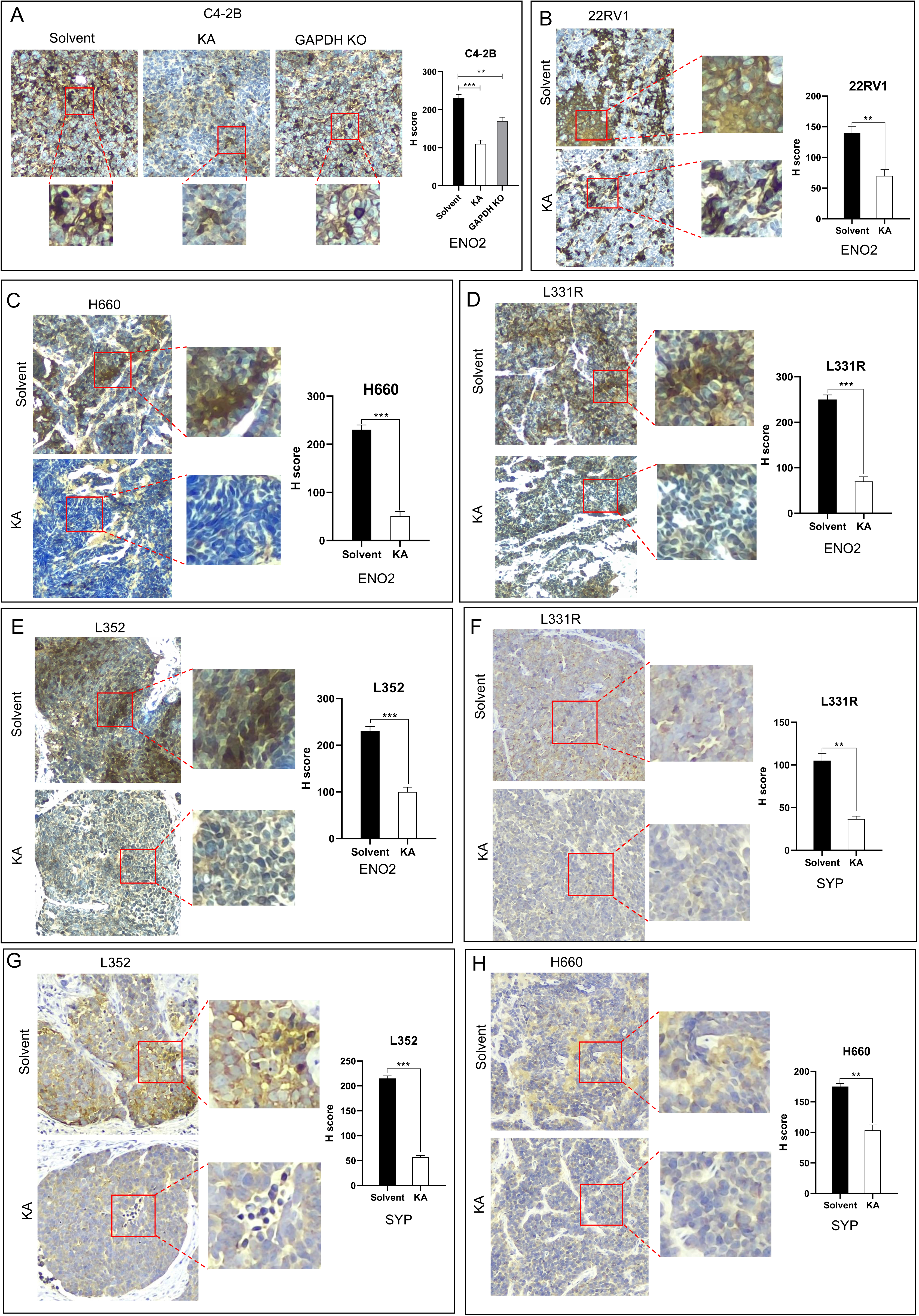
GAPDH inhibition suppresses neuroendocrine marker expression and tumor proliferative activity in CRPC and NEPC models. **A-E** ENO2 expression in CRPC and NEPC tumor tissues was evaluated by immunohistochemistry. **F-H** SYP expression in CRPC and NEPC tumor tissues was evaluated by immunohistochemistry.

**Fig 8.**
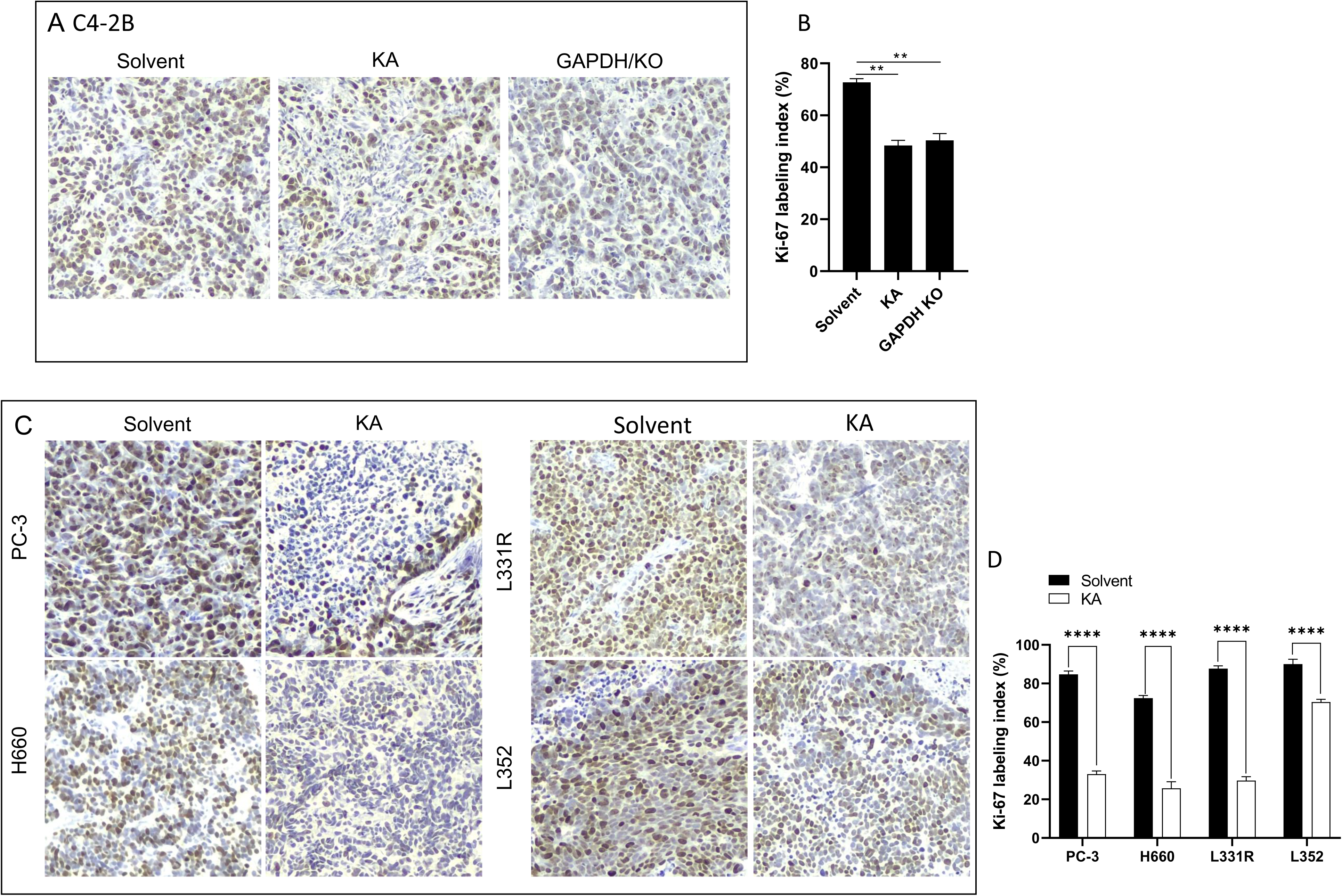
GAPDH inhibition suppresses cancer cell proliferation (Ki-67) in CRPC and NEPC tumors. **A, B** Parental and GAPDH/KO C4-2B xenograft tissue sections derived from solvent or KA treated groups were utilized for Ki-67 IHC. Semi-quantitative data of Ki-67 positive rates were summarized in panel B. **C, D** Ki-67 IHC for the tumor sections as indicated and semi-quantitative data.

### MZF1 mediates Enzalutamide-induced GAPDH upregulation

To elucidate the mechanism underlying AR antagonist-induced GAPDH upregulation, we used a luciferase reporter assay coupled with a pharmacological approach to determine the cellular signal pathways involved in Enzalutamide-induced GAPDH expression. The Renilla luciferase reporter construct (hGAPDH-rLUC) was driven by a 0.5-kb human GAPDH minimum promoter (41, 42). Under ADT conditions (Fig 9A), Enzalutamide treatment markedly increased GAPDH promoter activity (> 9-fold), indicating that regulatory elements within the 0.5-kb promoter region contribute to AR antagonist-induced GAPDH transcription.

**Fig 9.**
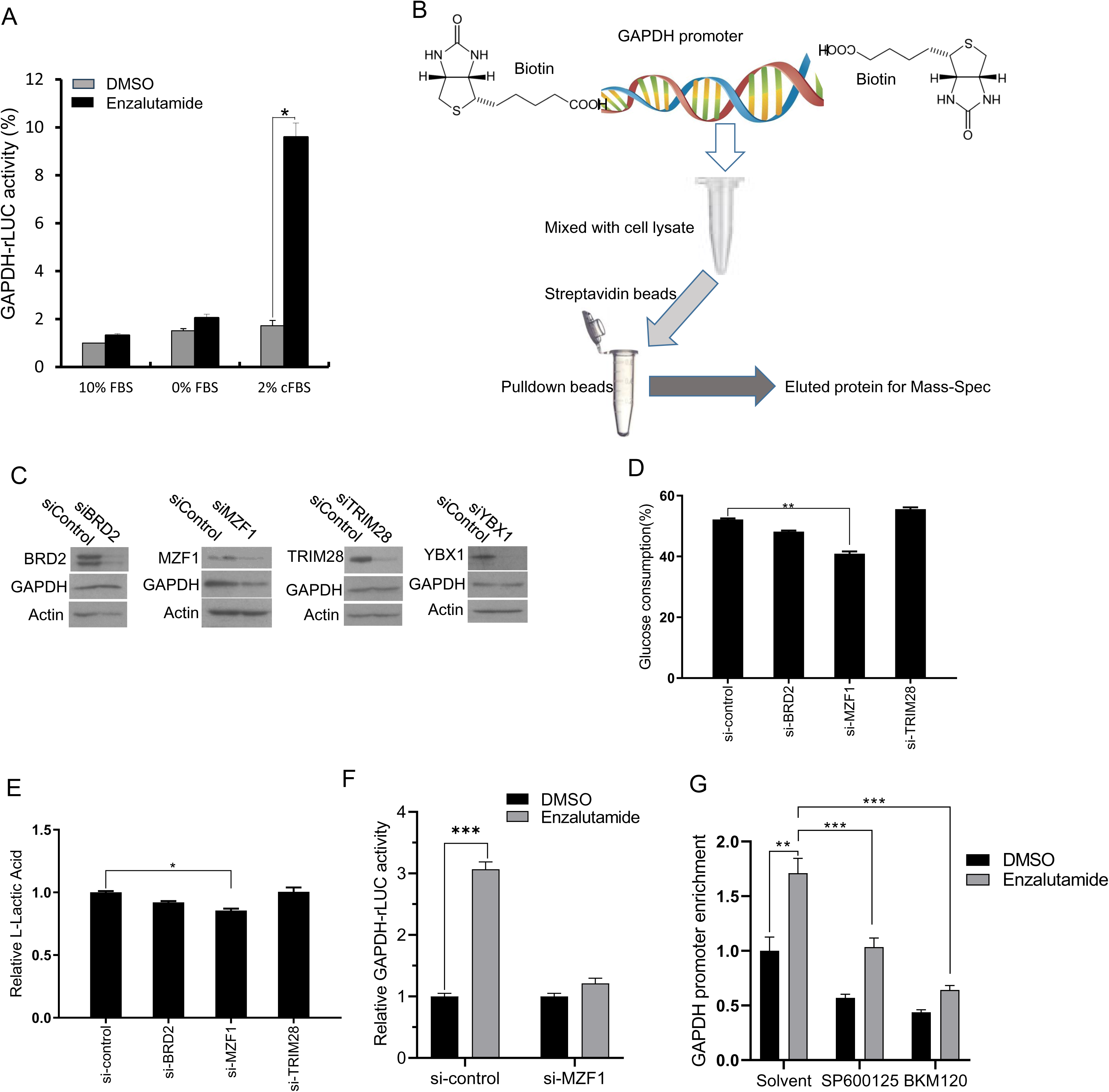
MZF1 mediates Enzalutamide-induced GAPDH transcriptional activation. **A** C4-2B cell was transfected with the hGAPDH-rLUC reporter plasmid and then treated with Enzalutamide (10 μM) overnight under 2% cFBS conditions. GAPDH promoter activity was measured by luciferase assay. **B** Schematic illustration of the GAPDH promoter DNA pulldown assay. **C** C4-2B cells were transfected with gene-specific ON-TARGETplus siRNA for 72 h, followed by immunoblot analysis of candidate GAPDH promoter-binding proteins and GAPDH. Actin served as a loading control. **D, E** C4-2B cells were transfected with the indicated siRNAs for 72 h, followed by medium replacement and overnight culture. Glucose consumption and L-lactate production were then measured. **F** C4-2B cells were treated with DMSO or Enzalutamide (10 μM) under 2% cFBS conditions and analyzed by anti-MZF1 ChIP assay. Enrichment of MZF1 at the GAPDH promoter was quantified by qPCR. RPL30 primers were used as an internal control. **G** C4-2B cells were transferred with the control or MZF1-specific siRNAs for 48 h, followed by transfection with the hGAPDH-rLUC reporter plasmid for 24 h. After overnight treatment by Enzalutamide (10 μM) with/without SP600125 (10 μM) or BKM120 (10 μM), cells were harvested for the luciferase assay. Data are presented as mean ± SEM. Asterisks indicate statistical significance compared with DMSO or Enzalutamide-treated controls; * p < 0.05, ** p < 0.01.

To identify the transcription factors involved in GAPDH promoter activation, we established a DNA-protein pulldown assay using the GAPDH promoter fragment (Fig 9B). Mass spectrometry analysis of eluted proteins recovered from the promoter-protein complex identified three candidate proteins with the highest confidence, including MZF1, BRD2, TRIM28, and YBX1 (Supplemental Fig S2, Supplemental Table S4).

To determine the functional relevance of these candidates, siRNA-mediated knockdown experiments were performed. Silencing MZF1, but not BRD2, TRIM28, and YBX1, significantly reduced GAPDH protein expression (Fig 9C). Furthermore, MZF1 gene silencing reduced glucose consumption, L-Lactate production, and hGAPDH-rLUC reporter activity in C4-2B cells (Fig 9D - 9F), supporting a role for MZF1 in regulating glycolysis and GAPDH transcriptional expression. To confirm the MZF1 interaction with the GAPDH gene promoter, CHIP assays were performed in C4-2B cells. Enzalutamide treatment significantly increased MZF1 occupancy at the GAPDH promoter region. Pretreatment with JNK and PI3K inhibitors significantly reduced MZF1 interaction with the GAPDH promoter, indicating both JNK and PI3K signal pathways are involved in modulating MZF1 transactivation towards GAPDH expression (Fig 9G).

Collectively, these findings verified MZF1 as a transcriptional mediator of Enzalutamide-induced GAPDH upregulation (Fig 10). These observations are consistent with previous reports showing MZF1-dependent regulation of GAPDH expression in multiple cellular contexts (43).

**Fig 10.**
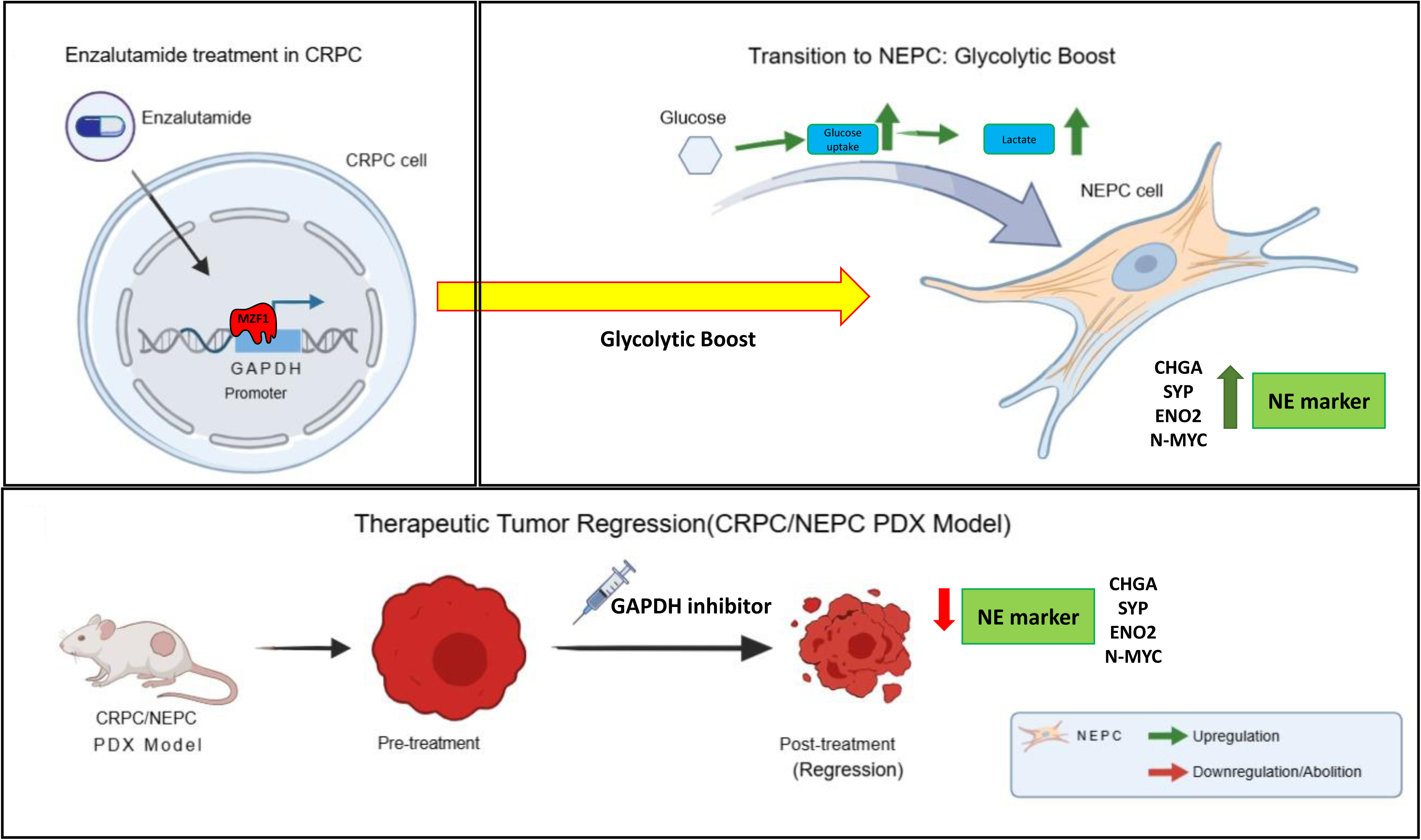
Schematic model summarizing the proposed mechanism by which AR antagonist treatment induces MZF1-dependent GAPDH activation, glycolytic reprogramming, and neuroendocrine-associated molecular adaptation, and how GAPDH inhibition suppresses this adaptive program.

## Discussion

Treatment-emergent neuroendocrine prostate cancer (t-NEPC) is increasingly recognized as a clinically aggressive form of prostate cancer that arises during prolonged androgen receptor (AR)-targeted therapy. Although lineage plasticity driven by transcriptional and epigenetic remodeling has been extensively implicated in this process (44–49) [reviewed in ref. (5, 9, 50)]. In parallel to these transcriptional and epigenetic alterations, metabolic reprogramming was also reported during neuroendocrine plasticity, as shown by our previous study (15) and recently by others (51–58). In the present study, we identified a previously unrecognized metabolic adaptation program induced by AR antagonists in CRPC cells. Our findings demonstrate that AR antagonist treatment induces glycolytic activation through MZF1-dependent transcriptional upregulation of GAPDH, leading to broad transcriptomic and metabolic remodeling, increased expression of neuroendocrine-associated genes, and enhanced tumor growth. Importantly, genetic or pharmacological inhibition of GAPDH suppressed glycolytic activation, attenuated neuroendocrine-associated molecular programs, and inhibited tumor growth in both CRPC and NEPC models, supporting GAPDH as a potentially actionable metabolic vulnerability during treatment-induced lineage plasticity.

Enhanced glycolysis is increasingly recognized as a hallmark of neuroendocrine plasticity in prostate cancer. Previous studies demonstrated dysregulation of multiple glycolytic components during NEPC development, including HK2/PFKP (55), PKLR (59), MCT4 (15), and ENO2 (56). Consistent with these observations, our study showed that AR antagonist treatment enhanced glucose uptake and glucose consumption under androgen-deprived conditions and markedly increased GAPDH expression and enzymatic activity. Interestingly, this metabolic response occurred not only in AR-positive CRPC models but also in AR-negative PC-3 cells, suggesting that AR antagonists may induce adaptive metabolic responses through mechanisms that are not strictly dependent on canonical AR signaling. These observations are consistent with the concept that therapeutic stress itself can drive metabolic reprogramming and cellular adaptation in prostate cancer.

Among the findings of this study, identification of MZF1 as a mediator of AR antagonist-induced GAPDH transcription provides a mechanistic link between AR-targeted therapy and glycolytic adaptation. Through promoter pulldown screening, mass spectrometry analysis, siRNA validation, and ChIP assays, we demonstrated that MZF1 directly contributes to GAPDH promoter activation following Enzalutamide treatment. Previous reports showed that MZF1 regulates glycolytic enzymes such as HK2 and phosphoglycerate kinase 1 (PGK1) in neuroblastoma cells (60), supporting a broader role for MZF1 in metabolic regulation. However, its role in treatment-induced metabolic adaptation in prostate cancer has not been previously defined. Notably, inhibition of Phosphoinositide 3-kinase (PI3K) and c-Jun N-terminal kinase (JNK) attenuated MZF1 transactivation towards GAPDH promoter interaction, suggesting that multiple cellular signal pathways may converge on MZF1-dependent transcriptional regulation during AR antagonist treatment. Nevertheless, the precise upstream mechanisms responsible for MZF1 activation remain to be elucidated. These observations support the concept that AR antagonist treatment itself may function as a metabolic stressor capable of triggering adaptive signaling and glycolytic reprogramming during therapeutic resistance.

An important observation from our study is that GAPDH inhibition not only reduced glycolytic activity but also suppressed neuroendocrine-associated molecular programs. GAPDH depletion or pharmacological inhibition reduced expression of canonical neuroendocrine markers, including CHGA, SYP, ENO2, and MYCN. In addition, transcriptomic analysis revealed widespread alterations in pathways associated with cholesterol biosynthesis, transcriptional regulation, central carbon metabolism, and peptidase-associated signaling following GAPDH depletion. Metabolomic profiling further demonstrated substantial metabolic remodeling characterized by the accumulation of glycolytic and TCA cycle intermediates and depletion of multiple amino acids. Together, these findings suggest that GAPDH-dependent glycolytic activation may contribute not only to metabolic adaptation but also to maintenance of neuroendocrine-associated cellular states during treatment-induced lineage plasticity.

Targeting glycolytic metabolism has emerged as a promising therapeutic strategy for overcoming treatment resistance in highly glycolytic tumors(17, 61–64). Because GAPDH occupies a central position in glycolytic flux, pharmacological inhibition of GAPDH may effectively disrupt adaptive metabolic programs associated with treatment resistance. Consistent with previous studies demonstrating antitumor activity of koningic acid (KA) in multiple cancer models (17, 19, 63–67), we observed significant tumor suppression in CRPC xenografts and NEPC PDX models following KA treatment. Importantly, KA suppressed tumor growth in models resistant to AR-targeted therapy, supporting the concept that metabolic targeting may bypass conventional resistance mechanisms. Interestingly, the LTL352 PDX model exhibited reduced sensitivity to KA and required a higher dose for therapeutic efficacy, suggesting heterogeneity in glycolytic dependency among NEPC subtypes. This observation further supports the notion that metabolic plasticity may differ substantially across treatment-emergent neuroendocrine states.

To further explore clinically relevant GAPDH-targeting approaches, we evaluated PGG as a reversible GAPDH inhibitor (40). PGG markedly suppressed LTL352 tumor growth with relatively limited systemic toxicity. Previous studies have also demonstrated antitumor activity of PGG in breast cancer (68), colon cancer (69), pancreatic cancer (70), and lung cancer (71), supporting its broader therapeutic potential. Collectively, these findings suggest that pharmacological targeting of GAPDH-dependent metabolic programs may represent a feasible therapeutic strategy for treatment-resistant prostate cancer, particularly in tumors exhibiting neuroendocrine-associated metabolic reprogramming.

Several limitations should also be acknowledged. Although our findings identify MZF1 as a critical mediator of GAPDH transcriptional activation, the upstream signaling events linking AR antagonist treatment to MZF1 activation remain incompletely understood. Furthermore, whether additional glycolytic enzymes cooperate with GAPDH during neuroendocrine plasticity requires further investigation. Because metabolic adaptation is highly dynamic, compensatory metabolic pathways may also contribute to therapeutic resistance following prolonged GAPDH inhibition. Finally, validation in additional translational and clinical settings will be required to determine the therapeutic applicability of GAPDH-targeted approaches in patients with treatment-emergent NEPC.

In conclusion, our study identifies an AR antagonist-MZF1-GAPDH signaling axis that promotes glycolytic activation, metabolic remodeling, and neuroendocrine-associated molecular reprogramming in prostate cancer. Targeting GAPDH suppressed tumor growth in CRPC and NEPC models and attenuated neuroendocrine-associated programs, supporting metabolic intervention as a potential therapeutic strategy for treatment-resistant prostate cancer (Fig 10).

## List of abbreviations

ADT: Androgen deprivation therapy
AR: Androgen receptor
ABI: abiraterone
APA: apalutamide
BSTFA: N-methyl-N-(trimethylsilyl) trifluoro-acetamide
CSDX: bicalutamide (Casodex)
CETSA: Cellular thermal shift assay
cFBS: Charcoal-filtered fetal bovine serum
CHGA: Chromogranin A
ChIP: Chromatin immunoprecipitation
CRPC: Castration-resistant prostate cancer
DAB: 3,3′-Diaminobenzidine
ENO2: Enolase 2
ENZA: Enzalutamide
GAPDH: Glyceraldehyde-3-phosphate dehydrogenase
GC-MS: Gas chromatography-mass spectrometry
GO: Gene Ontology
GSEA: Gene set enrichment analysis
HBSS: Hank’s Balanced Salt Solution
HF: hydroxy-flutamide
JNK: c-Jun N-terminal kinase
KA: Koningic acid
KO: Knockout
LC-MS/MS: Liquid chromatography-tandem mass spectrometry
MZF1: Myeloid zinc finger 1
MYCN / N-Myc: v-myc myelocytomatosis viral-related oncogene (neuroblastoma derived)
NEPC: Neuroendocrine prostate cancer
NF-κB: Nuclear factor kappa B
NSG: NOD-SCID-Gamma2 (mice)
PCA: Principal component analysis
PDX: Patient-derived xenograft
PGG: 1,2,3,4,6-penta-O-galloyl-β-D-glucopyranose
PI3K: Phosphoinositide 3-kinase
RIPA: Radioimmunoprecipitation assay
RNA-seq: RNA sequencing
STR: Short Tandem Repeat
SYP: Synaptophysin
TCA: Tricarboxylic acid
t-NEPC: Treatment-induced neuroendocrine prostate cancer

## Declarations

### Ethics approval and consent to participate

The study was conducted in accordance with the Declaration of Helsinki. All animal experimental protocols were approved by the Institutional Animal Care and Use Committee (IACUC) of the University of Kansas Medical Center (KUMC) (Protocol ID: 2020-2547-1). Patient-derived xenograft (PDX) models were maintained and utilized according to the ethical guidelines provided by the University of British Columbia and KUMC.

### Availability of data and materials

RNA-seq data were submitted to NCBI Bioproject website (#SUB16221405) pending approval. All data described in this study are available from the corresponding author upon request. All resource identifiers for antibodies and cell lines used in this study are provided in the Methods section.

### Authors’ contributions

WL and BL conceived and designed the study. WL and LH performed the experiments. DZ, CZ, and YW provided bioinformatic analysis. MM assisted with clinical data interpretation. WL and BL analyzed the data and wrote the manuscript. All authors read and approved the final manuscript.

## Acknowledgements

We thank the members of the Li Laboratory at the University of Kansas Medical Center for their technical assistance and helpful discussions throughout this study.

## Supplemental Materials

**Figure S1. AR antagonists fail to induce glycolytic activation in benign prostate cells.**

**A** BPH1 cells were treated with DMSO or the indicated AR antagonists for 24 h under 2% cFBS conditions. Glucose consumption was measured using a glucose oxidase-based assay and calculated as [(before − after)/before × 100%].

**B** Cells were treated with DMSO or Enzalutamide under 2% cFBS conditions for the indicated times, followed by GAPDH protein analysis.

**Fig S2. Identification of candidate GAPDH promoter-associated transcriptional regulators.** The Venny map for GAPDH promoter-binding proteins identified by DNA pulldown coupled with LC-MS/MS analysis.

Table S1, RNAseq data.

Table S2, GSEA-UPFC2 list.

Table S3, GAPDH/KO metabolomics.

Table S4, GAPDH promoter-binding proteins.

Table S5, Reagents and compounds for this study.

Table S6, Antibodies for WB and IHC.

Table S7, Primer sequences used for quantitative PCR.

