## Supplementary figures and images for "MZF1-mediated GAPDH overexpression drives glycolytic reprogramming and neuroendocrine progression in advanced prostate cancer"

### Supplemental Figure 1

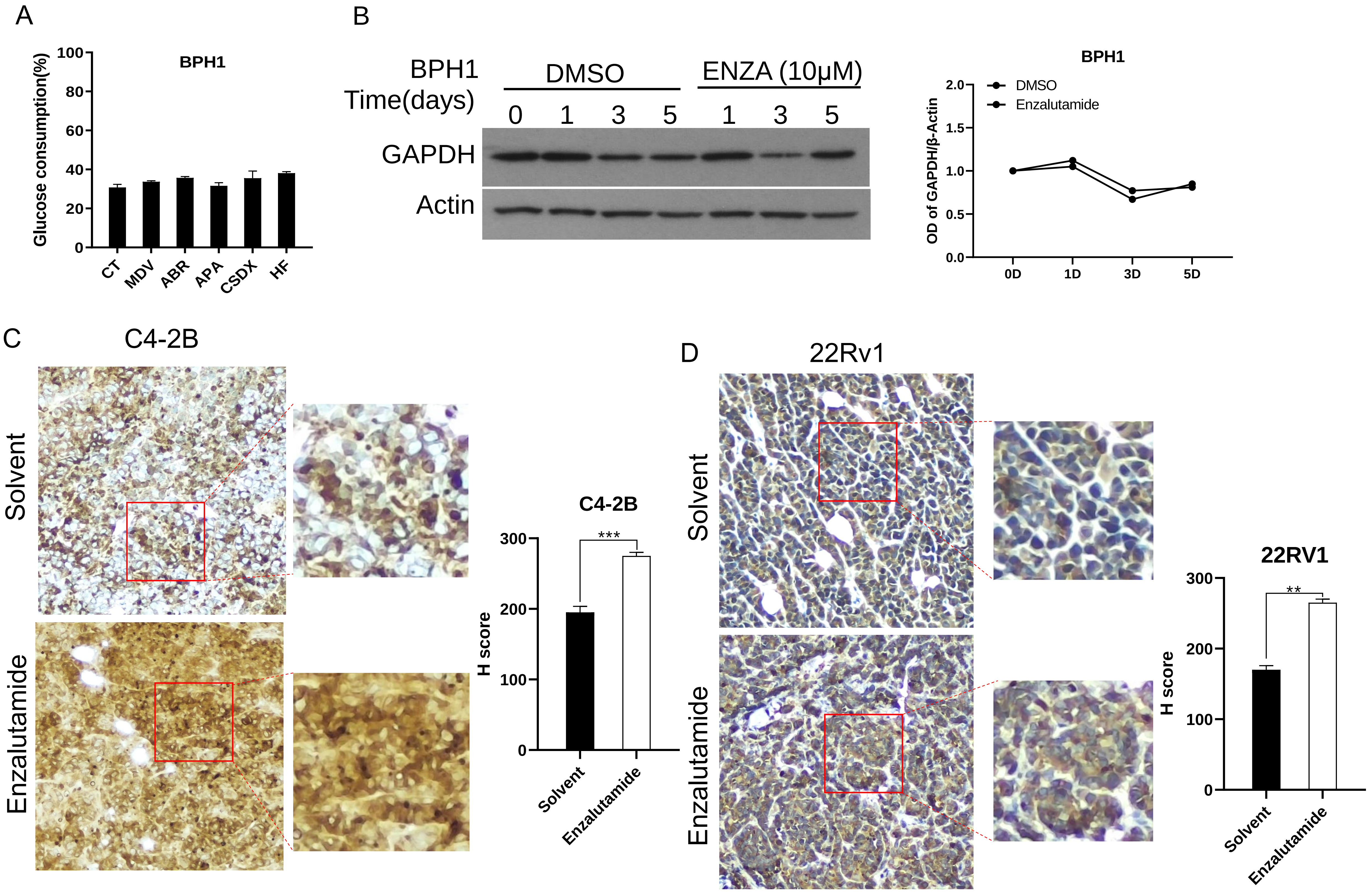

### Supplemental Figure 2

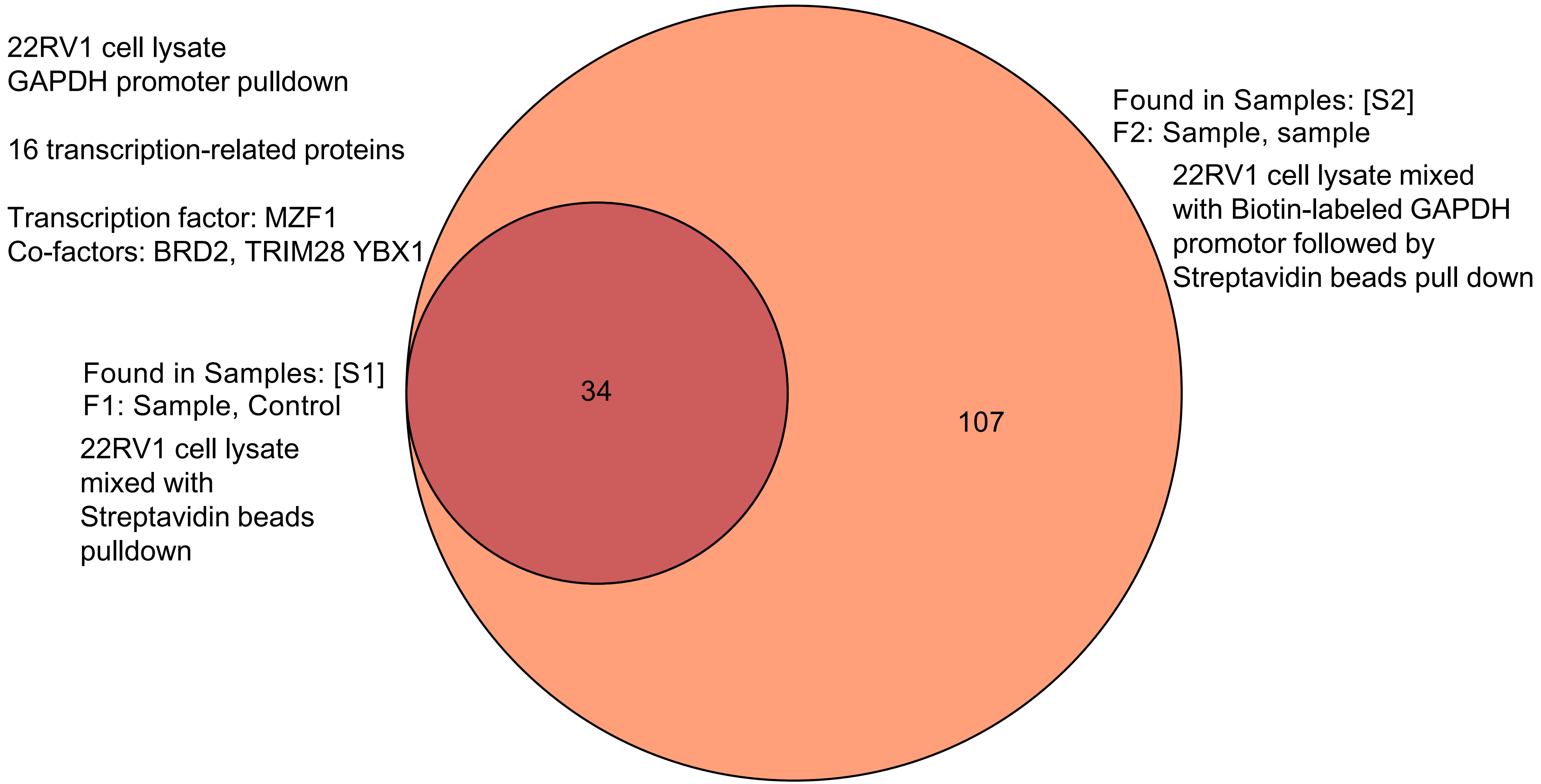
